# Dense cortical input to the rostromedial tegmental nucleus mediates aversive signaling

**DOI:** 10.1101/2021.09.10.459663

**Authors:** Elizabeth J Glover, E Margaret Starr, Andres Gascon, Kacey Clayton-Stiglbauer, Christen L Amegashie, Alyson H Selchick, Dylan T Vaughan, Wesley N Wayman, John J Woodward, L Judson Chandler

## Abstract

The rostromedial tegmental nucleus (RMTg) encodes negative reward prediction error (RPE) and plays an important role in guiding behavioral responding to aversive stimuli. While initial studies describing the RMTg revealed the presence of cortical afferents, the density and distribution of this input has not been explored in detail. In addition, the functional consequences of cortical modulation of RMTg signaling are only just beginning to be investigated. The current study anatomically and functionally characterizes cortical input to the RMTg in rats. Findings from this work reveal dense input spanning the entire medial prefrontal cortex (PFC) as well as the orbitofrontal cortex and anterior insular cortex. Afferents were most dense in the dorsomedial subregion of the PFC (dmPFC), an area which has also been implicated in both RPE signaling and aversive responding. RMTg-projecting dmPFC neurons originate in layer V and collateralize extensively throughout the brain. In-situ mRNA hybridization further revealed that neurons in this circuit are predominantly D1 receptor-expressing with a high degree of D2 receptor colocalization. Optogenetic stimulation of dmPFC terminals in the RMTg drives avoidance, and cFos expression is enhanced in this neural circuit during exposure to aversive stimuli. Exposure to such aversive stimuli results in significant physiological and structural plasticity suggestive of a loss of top-down modulation of RMTg-mediated signaling. Altogether, these data reveal the presence of a prominent cortico-subcortical projection involved in adaptive behavioral responding and provide a foundation for future work aimed at exploring alterations in circuit function in diseases characterized by deficits in cognitive control over the balance between reward and aversion.

## Introduction

Adaptive responding, in which past outcomes shape decision-making and future behaviors, is critical for survival. Impaired decision-making and maladaptive behaviors are common to a number of neuropsychiatric diseases and contribute significantly to harm and illness severity. Consequently, substantial effort has been put forth to identify the neural mechanisms governing such motivated behavior.

In the early 1980s, Corbett & Wise (1980) found that intracranial self-stimulation was most robust in rats with electrodes implanted in the ventral tegmental area (VTA) thereby uncovering a role for midbrain dopamine neurons in reward processing. Building upon this work, subsequent studies revealed that the activity of midbrain dopamine neurons encode a reward prediction error (RPE) with outcomes that are greater than expected producing a positive RPE associated with increased VTA dopamine neuron activity, and outcomes that are worse than expected producing a negative RPE associated with decreased VTA dopamine neuron activity (Schultz, 1986; Schultz et al., 1997). The neural circuitry driving RPE calculations within the VTA remains an area of intense investigation. To date, research suggests that positive RPE signals in the VTA are driven, at least in part, by excitatory input arising from the pedunculopontine tegmental nucleus (PPTg) (Mena-Segovia and Bolam, 2017) with possible additional involvement from neurochemically heterogenous afferents originating in the lateral hypothalamus (Nieh et al., 2015; Sharpe et al., 2017). Non-human primate studies suggested that the lateral habenula (LHb) played an important role in negative RPE signaling (Matsumoto and Hikosaka, 2007). However, this work presented somewhat of a paradox as previous studies had demonstrated that the LHb provides monosynaptic glutamatergic input to VTA dopamine neurons (Omelchenko et al., 2009) making it difficult to reconcile how this projection could facilitate a depression in phasic dopamine activity during presentation of aversive stimuli.

The solution to this paradox presented itself when two independent laboratories identified a previously unknown brain region called the rostromedial tegmental nucleus (RMTg), or tail of the VTA (Kaufling et al., 2009; Jhou et al., 2009b). Located immediately posterior to the VTA, both groups showed that the RMTg is primarily comprised of GABAergic neurons that receive dense input from the LHb and exert inhibitory control over monoaminergic and cholinergic midbrain nuclei including VTA dopamine neurons. Subsequent work found that RMTg activity increases in response to aversive stimuli of various sensory modalities (Jhou et al., 2009a; Li et al., 2019a) and that loss of RMTg function enhances active (e.g., escape) while reducing passive (e.g., freezing) responding in tests measuring fear and learned helplessness (Jhou et al., 2009a; Elmer et al., 2019). Circuit-specific approaches have further revealed that stimulation of dopamine-projecting RMTg neurons produces avoidance in a real-time place preference test and increases immobility in the forced swim test (St Laurent et al., 2020; Sun et al., 2020). In addition, using in-vivo electrophysiology, Hong et al. (2011) demonstrated that inhibition of VTA dopamine neurons in response to aversive stimuli is driven via activation of a disynaptic circuit comprised of glutamatergic LHb neurons that provide input to VTA-projecting GABAergic neurons in the RMTg. Collectively, these studies suggest that the LHb-RMTg-VTA circuit plays a critical role in encoding negative RPE thereby guiding behavioral responding to aversive stimuli.

The prefrontal cortex (PFC) integrates incoming multisensory information with previous experience to provide top-down inhibitory control over behavior and guide goal-directed responding. Interestingly, subregions spanning the dorsomedial PFC (dmPFC), which includes the anterior cingulate cortex (ACC) and prelimbic (PL) PFC in rats, corresponding to Broadman’s area 24 and 32 in humans, respectively, exhibit a number of functional similarities with the RMTg. For example, similar to the RMTg, neuronal activity in the PL PFC increases during exposure to aversive stimuli (Burgos-Robles et al., 2009) and loss of PL function reduces passive fear responding (Corcoran and Quirk, 2007). In addition, the activity in the dmPFC, and ACC in particular, has been heavily implicated in RPE signaling (Alexander and Brown, 2019). As with the dmPFC, the rodent ventromedial PFC (vmPFC) is also comprised of two subregions. The more ventral rodent infralimbic (IL) PFC, which is thought to be homologous with Broadman’s area 25 in humans, is well-characterized as frequently exerting opposing functions to the more dorsal PL mPFC (Peters et al., 2009; Gourley and Taylor, 2016) with activity in this region facilitating extinction of responses driven by PL PFC activity (Do-Monte et al., 2015). The dorsal peduncular cortex (DP) makes up the ventral-most portion of the vmPFC. While some have suggested the DP exhibits functional overlap with the IL PFC (Peters et al., 2009), very few studies have examined the DP directly and the human homolog of this region is, to date, unknown.

It is of interest that initial anatomical characterization of the RMTg revealed the presence of cortical afferents to the RMTg, including some that arose from the mPFC (Kaufling et al., 2009; Jhou et al., 2009b). However, the anatomy and function of these inputs have not been well-characterized as much of the research aimed at investigating the role of RMTg-associated neural circuits in aversive signaling has focused on the LHb-RMTg-VTA projection. The present study begins to fill this gap by anatomically and functionally characterizing cortical inputs to the RMTg with particular focus on those arising from the dmPFC given the functional overlap it shares with the RMTg. Our findings reveal the presence of dense input spanning a number of functionally distinct regions of the prefrontal and insular cortices and uncover a role for RMTg-projecting dmPFC neurons in top-down control over RMTg-mediated aversive signaling.

## Materials & Methods

### Animals

For all experiments, adult male Long-Evans rats (P60 upon arrival, Envigo Laboratories, Indianapolis, IN) were individually housed in standard polycarbonate cages. The vivarium was maintained on a 12:12 reverse light-dark cycle with lights off at 09:00. Rats were habituated to the vivarium for at least one week before beginning experiments. All rats were provided with Teklad 2918 (Envigo) standard chow and water *ad libitum*. All experiments were approved by the Institutional Animal Care and Use Committees at the Medical University of South Carolina and University of Illinois at Chicago and adhered to the guidelines put forth by the NIH (National Research Council, 2011).

### Stereotaxic surgery

Rats undergoing stereotaxic surgery were induced and maintained under a surgical plane of anesthesia using isoflurane (1-5%). Intracranial injections were made with back-filled custom glass pipettes connected to a nanojector (Drummond Scientific Company, Broomall, PA, USA) or a 200 uL Hamilton syringe operated using a motorized pump (WPI, Inc, Sarasota, FL, USA). For tract tracing experiments, 100 nL 0.5% Cholera toxin B (CtB; Sigma Aldrich, St Louis, MO, USA) or 200 nL green fluorescent retrobeads (Lumafluor, Durham, NC, USA) were unilaterally injected into the RMTg (AP: −7.3; ML: +1.4; DV: −8.0 from skull; 6° lateral) at a rate of ~30 nL/s. For optogenetics experiments, rats were bilaterally injected with 500 nL of AAV2-hSyn-hChR2-(H134R)-eYFP-WPRE-pA (UNC Vector Core, Chapel Hill, NC, USA) into the dmPFC (AP: +3.2; ML: ±0.6; DV: −3.5 from skull) or lateral habenula (LHb; AP: −3.6; ML: ±1.2; DV: −4.1 from dura; 6° lateral) at a rate of 1-3 nL/s. During the same surgery, rats were implanted with custom-made 200 μm optic fiber implants targeting either the RMTg (AP: −7.3; ML: ±2.1; DV: −7.9 from skull; 10° lateral) or VTA (AP: −5.6; ML: ±1.4; DV: −7.8 from skull; 6° lateral). Implants were secured with dental cement. An intersectional, dual-virus approach was used to investigate the extent of dmPFC-RMTg collateralization and structural plasticity in dmPFC-RMTg neurons following exposure to aversive stimuli. For these experiments, rats were unilaterally injected with 500 nL of either AAVretro-Cre (gift from Janelia Farms, Ashburn VA, US) or AAV2retro-pmSyn1-EBFP-Cre (Addgene, Watertown, MA, USA) into the RMTg (AP: −7.3; ML: +1.4; DV: −8.0 from skull; 6° lateral) and 500 nL of AAV8.2-hEF1alpha-DIO-SYP-EYFP (Rachel Neve, MIT Vector Core) into the dmPFC (AP: +3.2; ML: +0.6; DV: −3.5 from skull) at a rate of 1-3 nL/s.

### Cell density analysis

Rats unilaterally injected with CtB into the RMTg (n=9) were transcardially perfused with phosphate buffered saline (PBS) followed by 4% paraformaldehyde (PFA). Brains were immersion fixed overnight in 4% PFA, cryoprotected in 30% sucrose, and stored at −80 °C until ready for processing for microscopic analysis. Brains were sliced at 40 μm on a cryostat held at - 20 °C. Slices containing the PFC and RMTg were labeled for CtB and NeuN using standard immunofluorescence procedures. In brief, slices were incubated in 50% (v/v) methanol for 30 min followed by incubation in 1% H_2_O_2_. Permeabilization was enhanced by incubation in 0.4% Triton-X in PBS followed by incubation in primary antibodies in PBS containing 0.2% Triton-X overnight at 4 °C (CtB 1°: 1:500, List Biological Laboratories #703; NeuN 1°: 1:500, EMD Millipore, MAB377). The tissue was then incubated with secondary antibodies for 2 h at room temperature (1:250, Jackson ImmunoResearch), rinsed in PBS, and mounted onto SuperFrost plus charged slides before being coverslipped with Fluoromount mounting medium (Sigma Aldrich). Images were acquired at 10X and tiled using an EVOS FL Auto microscope. Anatomical boundaries and rostrocaudal level of each PFC slice were determined by aligning the acquired microscopic images with atlas schematics generated using Paxinos & Watson (2007) in GIMP. ImageJ was used to apply a bandpass filter to Fourier-transformed images after which CtB+ and NeuN+ cells were automatically identified by searching for maximum intensity points.

### Cell-type analysis

Adjacent tissue from that used in the cell density analysis experiments (n=3) was used to evaluate whether RMTg-projecting cortical neurons were glutamatergic or GABAergic projection neurons. Slices containing the PFC were labeled for CtB and the glutamatergic marker, CaMKIIα, or CtB and the GABAergic marker, GAD67, using the same immunofluorescence procedures described above (CaMKIIα 1°: 1:3,000, Invitrogen MA1048; GAD67 1°: 1:3,000, EMD Millipore MAB5406). Images of labeling in the dmPFC were acquired at 10X using a Zeiss AxioImager.M2 microscope. CtB-, CaMKIIα-, and GAD67-labeled cell bodies were counted manually in each image using ImageJ. Cell counts were averaged across five slices spanning the rostrocaudal extent of the dmPFC for each rat and the ratio of cells labeled with each cell-type marker and CtB relative to all CtB-labeled neurons was calculated.

### Collaterals analysis

A dual-virus, intersectional approach was used to label RMTg-projecting dmPFC neurons. After waiting at least eight weeks for optimal viral transduction and transgene expression, rats were euthanized and brains harvested using the same procedures described above. Brains were sliced on a cryostat at 40 μm and eYFP signal was amplified using avidin-biotin immunohistochemistry procedures as previously published (Glover et al., 2016; GFP Abcam, ab290; 1:10,000). Brains were visually inspected from the rostral tip of the PFC to the rostral cerebellum for eYFP+ terminal labeling. Areas with noticeable labeling were imaged at 10X on a Zeiss AxioImager.M2 microscope. Images were flat field corrected and terminal density was analyzed by measuring the percent-stained area relative to total area using ImageJ. Analysis was performed on four slices spanning the rostrocaudal extent of each region and averaged together to arrive at a single data point for each region. Analysis of secondary somatosensory cortex and dorsal hippocampus were included as negative controls.

### Real-time place preference testing

After at least eight weeks to allow for sufficient viral transduction and transgene expression, rats were tested for real-time place preference using procedures adapted from previously published work (Stamatakis and Stuber, 2012). Rats were habituated to the tethering procedure for at least three days prior to testing. Testing was performed in an unbiased, custom-made apparatus consisting of two contextually distinct compartments. On day one, rats were connected to a patch cable connected to a 473 nm laser and allowed to freely explore the apparatus for 20 min. Light was delivered immediately upon entry into one compartment of the apparatus at 10 mW intensity and 60 Hz for the duration of time spent in that compartment. Light delivery was terminated upon entry into the opposite compartment. To confirm that behavioral responding was light-mediated, rats were re-tested 24 hours later using identical procedures except that the compartment associated with light delivery was reversed. Time spent in each compartment was quantified from video recordings made with a camera mounted above the testing apparatus using Ethovision (Noldus, Leesburg, VA, USA).

### cFos induction following aversive stimuli

cFos induction was measured in RMTg-projecting mPFC neurons following exposure to aversive stimuli using procedures adapted from Jhou et al. (2009a). Rats were allowed at least one week to recover following stereotaxic injection of CtB into the RMTg before beginning testing. All rats underwent three days of habituation during which they were tethered and could freely explore a standard operant testing apparatus (Med Associates, St Albans, VT, USA). The house light was illuminated for the duration of each habituation session and all subsequent sessions. On day four, rats in the Context (control) group were euthanized 90 min after an identical 20 min habituation session. Rats in the Shock group were euthanized 90 min after presentation of a series of 10 foot shocks (0.5 mA, 0.5 s duration, 60 s inter-stimulus interval) over the course of a 20 min testing session beginning 60 s after the start of the session. Rats in the Shock-paired group underwent standard fear conditioning over two consecutive days where tone (2.9 kHz, 65 dB, 20 s duration) presentation co-terminated with foot shock (0.5 mA, 0.5 s duration). Two tone-shock pairings were presented 20 min apart over the course of each 60 min conditioning session. Rats in the Shock-unpaired group were presented with the same stimuli as the Shock-paired group during two 60 min sessions except that stimuli were explicitly unpaired occurring 10 min apart. Following conditioning trials, rats from both the Shock-paired and Shock-unpaired groups were re-habituated to the testing apparatus during a 30 min session where the house light was illuminated but no stimuli were presented. On the test day, rats from both groups were euthanized 90 min after a 20 min test session consisting of presentation of eight tones for 30 s each (60 s inter-stimulus interval). Freezing during tone presentation was scored manually in the Shock-paired and Shock-unpaired rats using overhead video recorded during the test session. Following euthanasia, brains were processed for CtB (1:300,000) and cFos (Millipore #PC38, 1:10,000) expression using previously published procedures (Glover et al., 2016). The number of double-labeled neurons relative to all CtB+ neurons was quantified manually across 4-5 slices spanning the rostrocaudal extent of the mPFC. Rats with off-target injection sites were excluded from analysis.

### In-situ hybridization

Rats were unilaterally injected with green retrobeads into the RMTg and allowed at least seven days to recover before testing. The animals were assigned to either Context or Shock groups and underwent testing identical to that described above for the cFos induction experiments. Rats were anesthetized with isoflurane and decapitated 90 min after the test session. The brains were then rapidly removed and placed in ice-cold PBS for ~5 minutes before being embedded in Tissue-Tek OCT media (Sukura Finetek Inc, Torrance, CA, USA) in Peel-A-Way cryo-embedding molds (Polysciences, Inc, Warrington, PA, USA) and covered with dry ice. The frozen tissue block was then extracted from the mold, wrapped in aluminum foil, and stored at −80 °C. Subsequently, 20 μm thick slices from the fresh-frozen brains were cut on a cryostat, mounted on SuperFrost Plus slides (Fisher Scientific, Hampton, NH), and stored at −80 °C until their use in in-situ hybridization experiments.

Fluorescence in-situ hybridization was performed using an Advanced Cell Diagnostics (ACD, Newark, CA) Multiplex RNAScope kit (catalog # 323100). RNA probes for dopamine D1 receptors (catalog # 317031), dopamine D2 receptors (catalog # 315641-C2), and cFos (catalog # 403591-C4) were also obtained from ACD. The RNAScope procedure was carried out according to the manufacturer’s instructions (available for download at www.acdbio.com) with the exception that the protease digestion step was omitted. We observed that omission of this step not only improved the fluorescence intensity of the mRNA transcripts (visually observed as punctate dots), but was also required for preservation of the fluorescent intensity of the green (alexa-488) retrobeads. For multiplex hybridization of D1 and D2 mRNA transcripts, the probes were labeled with Cy3 and Cy5, respectively. For multiplex hybridization of cfos and D1 mRNA transcripts, the probes were labeled with Cy3 and Cy5, respectively. Images were acquired on a Zeiss LSM880 confocal microscope across three PFC slices (5 images/slice) using a 63X oil objective. Imaging was restricted to areas of the dmPFC that exhibited retrograde bead labeling, which was mainly observed in cortical layer V. Quantification and colocalization of mRNA transcript dot within cells was performed on the captured images using Imaris Software (Bitplane, Zurich, Switzerland) following a previously published method (Centanni et al., 2019). This analysis utilized the Cell Module of Imaris, and can be summarized as follows: 1) Define and interactively threshold the cell nucleus based on DAPI staining (we used a minimum size of 5 μm and a filter of 0.5); 2) Define and interactively threshold the cell body based upon the DAPI identified nucleus in step 1; 3) Define and interactively threshold the mRNA transcript dots for each probe and for the retrobeads (we used a minimum size of 1 μm for both the transcript dots and retrobeads); 4) Calculation of the number and other parameters of the dots that lie within each defined cell. For a cell to be considered as positive for fluorescent beads or D1/D2 mRNA transcripts, it had to contain two or more dots/beads. We observed that a number of cells exhibited a variable level of background cfos mRNA irrespective of whether they were in the Control or Shock group. Therefore, for the purpose of assessment of the effect of shock on cfos mRNA expression, we used a threshold of 15 or more cfos transcript dots in order to consider a cell as being cfos+. This threshold was determined based on a comparison of the distribution of the cfos mRNA transcript dots in the control versus shocked conditions.

### Whole-cell patch-clamp slice electrophysiology

Rats were unilaterally injected with green retrobeads into the RMTg and allowed at least three days to recover before being assigned to either Context or Shock groups and undergoing testing identical to that described above for cFos induction experiments. Twenty-four hours following the final day of testing, the intrinsic excitability of dmPFC pyramidal neurons was determined using previously published procedures (Wayman and Woodward, 2018). In brief, rats were anesthetized with urethane (3.0mg/kg, i.p.) and perfused with an ice-cold sectioning solution consisting of (in mM): 200 sucrose, 1.9 KCl, 6 MgSO_4_, 1.4 NaH_2_PO_4_, 25 NaHCO_3_, 0.5 CaCl_2_, 10 glucose, and 0.4 ascorbic acid; pH 7.35-7.45 with 310-320 mOsm. The brains were then immediately harvested and coronal brain sections (300 μm) containing the dmPFC were sliced on a Leica VT1000S vibratome (Leica Biosystems, Buffalo Grove, IL) in oxygenated (95% O_2_; 5% CO_2_) sectioning solution and then transferred to a holding chamber containing normal artificial cerebrospinal fluid (aCSF; in mM): 125 NaCl, 2.5 KCl, 25 NaHCO_3_, 1.4 NaH_2_PO_4_, 1.3 MgCl_2_, 2 CaCl_2_, and 10 glucose; pH 7.35-7.45 with 310-320 mOsm. Brain slices were incubated at 34 **°**C for 30 minutes and allowed to recover at room temperature for an additional 45 minutes.

For current clamp recordings, brain slices were transferred to the recording chamber and perfused with oxygenated and heated (~34 °C) aCSF at a flow rate of 2 mL/min. The temperature was maintained during the course of the recordings with in-line and bath heaters (Warner Instruments, Hamden, CT). Retrobead-labeled layer V neurons within the dmPFC were visually identified using a Zeiss FS2 microscope (Zeiss, Thorndale, NY). Recording pipettes were constructed from thin-walled borosilicate capillary glass tubing (I.D.=1.17mm, O.D. 1.50mm; Warner Instruments, Hamden, CT), pulled with a horizontal pipette puller (P-97 Sutter Instrument Co., Novata, CA). Pipettes were filled with an internal solution containing (in mM): 120 K-gluconate, 10 HEPES, 10 KCl, 2 MgCl2, 2 Na2ATP, 0.3 NaGTP, 1 EGTA and 0.2% biocytin; pH 7.35-7.45 with 285-295 mOsm and had resistances ranging from 3-5 MΩ. After a stable gigaohm seal was formed, light suction was applied to break through the cell membrane and achieve whole-cell access. Neurons with an access resistance of greater than 20 mΩ were not used for analysis. Recorded events were acquired with an Axon MultiClamp 700A amplifier (Molecular Devices, Union City, CA), digitized at a sampling rate of 10 kHz (filtered at 4 kHz) with an Instrutech ITC-18 analog-digital converter (HEKA Instruments, Bellmore, NY) controlled by AxographX software (Axograph Scientific, Sydney, Australia) running on a Macintosh G4 computer (Apple, Cupertino, CA). The resting membrane potential (RMP) and capacitance of all neurons was first recorded and then the RMP was adjusted to −70 mV for electrophysiological assessments of excitability. Action potential firing was induced by a series of 500 ms current steps (0-300 pA) incremented in +20 pA steps. Recordings were analyzed offline for the number of spikes in response to each current step, threshold (mV), rheobase (pA), action potential peak amplitude (mV), action potential half-width (ms) and after-hyperpolarization (AHP; mV) using AxographX software.

The caudal portion of the brain containing the RMTg was collected at the same time that dmPFC slices were generated, immersion fixed overnight in 4% PFA, and frozen on dry ice followed by storage at −80 °C until processing. Injection sites were confirmed by visual inspection of fluorescent retrobead labeling in slices containing the RMTg made using a cryostat.

### Spine density analysis

A dual-virus, intersectional approach was used to label RMTg-projecting dmPFC neurons. After waiting at least eight weeks for optimal viral transduction and transgene expression, rats were assigned to either Context or Shock groups and underwent behavioral testing identical to that described above for cFos induction experiments. Twenty-four hours later, rats were euthanized, and brains harvested as described for cell density analysis. Brains were sliced at 100 μm and eYFP signal was amplified (GFP, Abcam #ab290; 1:30,000) using immunofluorescence procedures optimized for thick slices (Kupferschmidt et al., 2015). Primary apical dendrites measuring 55 μm in length approximately 200-300 μm from the soma of eYFP+ neurons in the dmPFC were imaged using a 63.5X oil immersion objective on a Zeiss LSM880 confocal microscope. Images were analyzed in Imaris using previously published procedures (McGuier et al., 2015). Dendrite diameter, dendrite volume, and total spine density were analyzed in addition to analyses conducted by spine classification. Measures included density, length, diameter, and volume by spine class as well as diameter and volume of spine terminal point and spine neck volume, length, and diameter. Measures were collected in 3-5 dendrites per rat and averaged across dendrites to arrive at a single value for each rat.

### Statistical analysis

Student’s t-test and analysis of variance (ANOVA) were performed to analyze all functional data as indicated below in the Results section. Greenhouse-Geisser correction was performed on data that lacked sphericity. All analyses were performed using GraphPad Prism 8.0 and are presented as mean ± SEM. Effects were considered statistically significant at p ≤ 0.05.

## Results

### Cortical input to the RMTg is dense, glutamatergic, and exhibits extensive collaterals

While initial reports indicated the presence of cortical efferents to the RMTg, the magnitude of this input and subregional distribution was unclear. To investigate this, RMTg-projecting cell bodies were quantified in rat brains injected with the retrograde tracer CtB. Visual inspection of slices double stained for CtB and the neuronal marker, NeuN, revealed the presence of dense input spanning the medial wall of the PFC and the entire OFC (**Figure 1A**). In line with previous reports, relatively low but consistent labeling was also observed in the anterior insular cortex (AIC) (Kaufling et al., 2009; Jhou et al., 2009b). In agreement with the well-understood layer specificity of cortico-subcortical projections in the rodent mPFC, the majority of CtB+ cell bodies originated in layer V (**Figure 1B**). Consistent cell body labeling was also apparent in the deepest portion of layer VI, albeit to a much smaller degree than was observed in layer V. Quantification of layer V CtB+ neurons relative to NeuN+ neurons in the mPFC revealed relatively uniform densities of RMTg-projecting neurons across the rostrocaudal extent of ACC (3.96 ± 0.28%), PL (8.59 ± 0.45%), and IL (7.95 ± 0.77%) subregions (**Figure 1C**). In contrast, the density of RMTg-projecting dorsopeduncular (DP) mPFC neurons increased substantially at more caudal levels relative to rostral DP mPFC (13.02 ± 2.91%).

**Figure 1.**
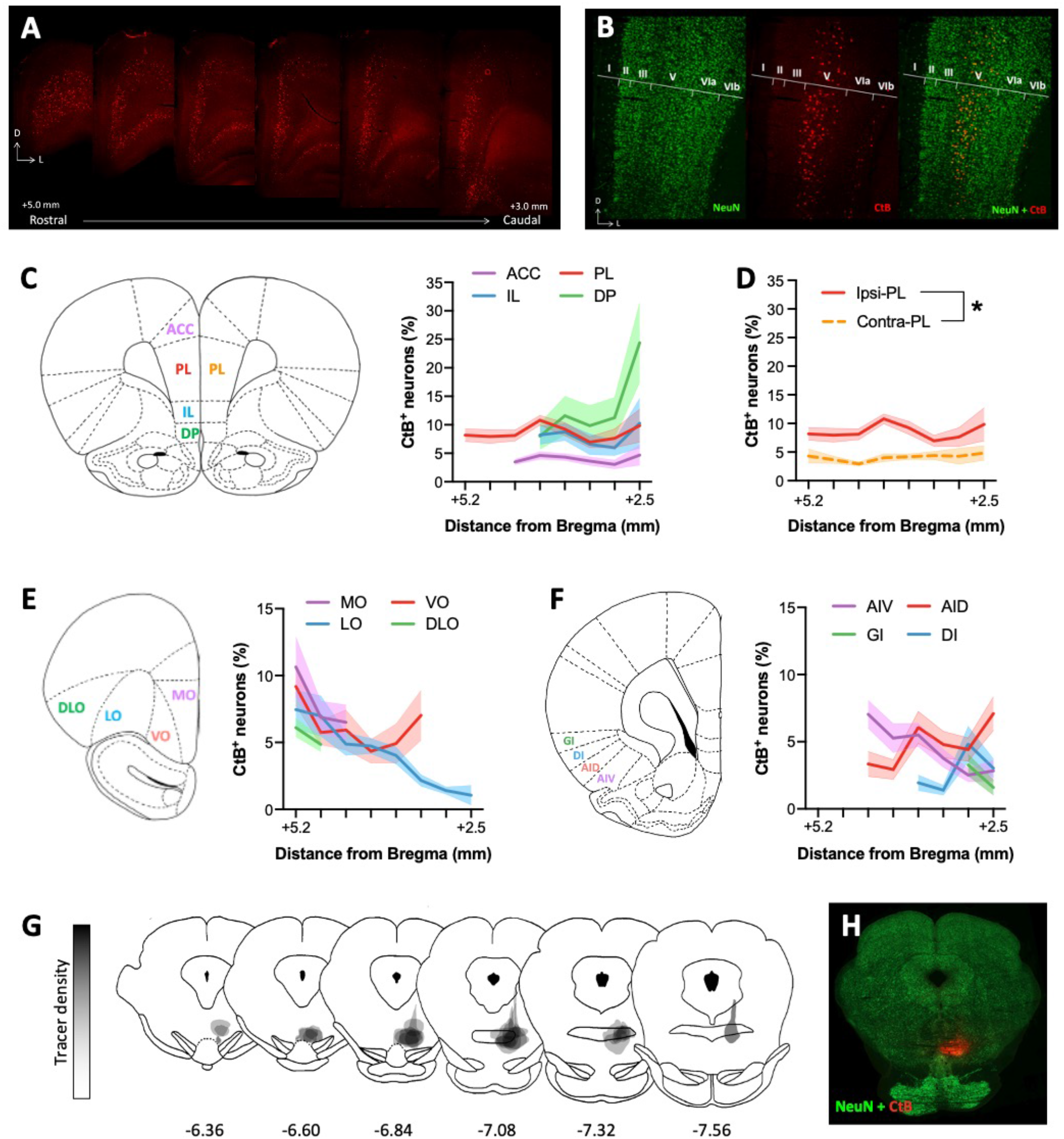
Anatomical distribution of cortical inputs to the RMTg. **(A)** Representative images demonstrating dense ipsilateral cortical labeling in brain areas injected with CtB into the RMTg. **(B)** Representative high magnification image showing that inputs to the RMTg arise primarily from layer V of the mPFC. **(C)** The percent of CtB+ neurons relative to all layer V NeuN+ neurons is relatively consistent across ACC, PL, and IL subregions of the mPFC whereas the density of RMTg-projecting DP mPFC neurons increases substantially at more caudal levels. **(D)** Contralateral cortical afferents are substantially less dense than ipsilateral inputs as exemplified by a comparison of RMTg-projecting PL mPFC neurons in both hemispheres. **(E)** The density of layer V OFC neurons projecting to the RMTg is similar to that observed in the mPFC with LO inputs diminishing at more caudal levels. **(F)** CtB labeling is consistently observed in the AIC, albeit to a lesser degree than that observed in mPFC and OFC. **(G)** Map of tracer injection sites for all animals included in quantification. **(H)** Representative injection site. Abbreviations: ACC = anterior cingulate cortex; AID = agranular insular cortex, dorsal; AIV = agranular insular cortex, ventral; DI = dysgranular insular cortex; DLO = dorsolateral orbitofrontal cortex; DP = dorsopeduncular cortex; GI = granular insular cortex; IL = infralimbic cortex; LO = lateral orbitofrontal cortex; MO = medial orbitofrontal cortex; PL = prelimbic cortex; VO = vental orbitofrontal cortex.

The density of CtB-labeled neurons was relatively similar across subregions of the OFC at rostral levels but began to diverge slightly more caudally (**Figure 1E**). On average, density was greatest and somewhat variable in the medial orbital (MO) cortex (8.01 ± 1.32%). By contrast, CtB labeling was lower and less variable in the dorsolateral orbital (DLO) cortex (5.48 ± 0.62%). The density of RMTg-projecting ventro-orbital (VO) and latero-orbital (LO) cortical projections varied substantially from rostral to more caudal levels within the brain. In the VO, density was greatest at the rostral-most point of the OFC (9.18 ± 1.43%) after which CtB labeling diminished (4.35 ± 0.98%) before increasing in density at its most caudal point (7.03 ± 1.87%). By contrast, RMTg-projecting neurons arising from the LO cortex are most dense at the rostral tip of the region (7.45 ± 1.43%) and become progressively less dense as one moves caudally with very little CtB+ cell bodies in the most caudal region (1.05 ± 0.73%).

Although substantially less dense than projections arising from the mPFC and OFC, CtB+ cell bodies were consistently observed in subregions of the AIC (**Figure 1F**). Density was greatest in the agranular AIC with approximately 4.5% of layer V neurons in dorsal (AID) and ventral (AIV) subregions projecting to the RMTg (AID: 4.77 ± 0.65%; AIV: 4.49 ± 0.72%). By contrast, CtB labeling was approximately half that of agranular AIC subregions in the dysgranular (DI; 2.8 ± 0.76%) and granular (GI; 2.43 ± 0.84%) cortices.

As is often the case, CtB labeling was most dense in the hemisphere ipsilateral to the injection site with substantially less labeling apparent in the contralateral cortex. To measure this directly, we quantified the density of layer V RMTg-projecting neurons in the contralateral PL mPFC and found that contralateral cell density was approximately half that of the ipsilateral projection (4.07 ± 0.20%) (**Figure 1D**). A two-way RM ANOVA comparing cell density between ipsilateral and contralateral hemispheres across the rostrocaudal extent of the PL mPFC confirmed that significantly fewer RMTg-projecting cells arise in the contralateral compared to ipsilateral hemisphere regardless of rostrocaudal level [main effect of hemisphere: F(1,16)=45.09, p<0.0001).

Layer V cortical efferents are typically excitatory in nature, however, recent work has revealed the presence of long-range GABAergic projection neurons arising from various cortical regions including the mPFC (Lee et al., 2014; Basu et al., 2016; Rock et al., 2018). To determine the neurochemical composition of RMTg-projecting cortical neurons, CtB-expressing slices adjacent to those used in the cell density analysis were labeled with CaMKIIα or GAD67. As shown in **Figure 2**, CtB-labeled neurons were predominantly CaMKIIα+. In contrast, virtually no overlap in expression was observed between CtB and the GABAergic marker, GAD67. These data indicate that RMTg-projecting cortical neurons are excitatory projection neurons.

**Figure 2.**
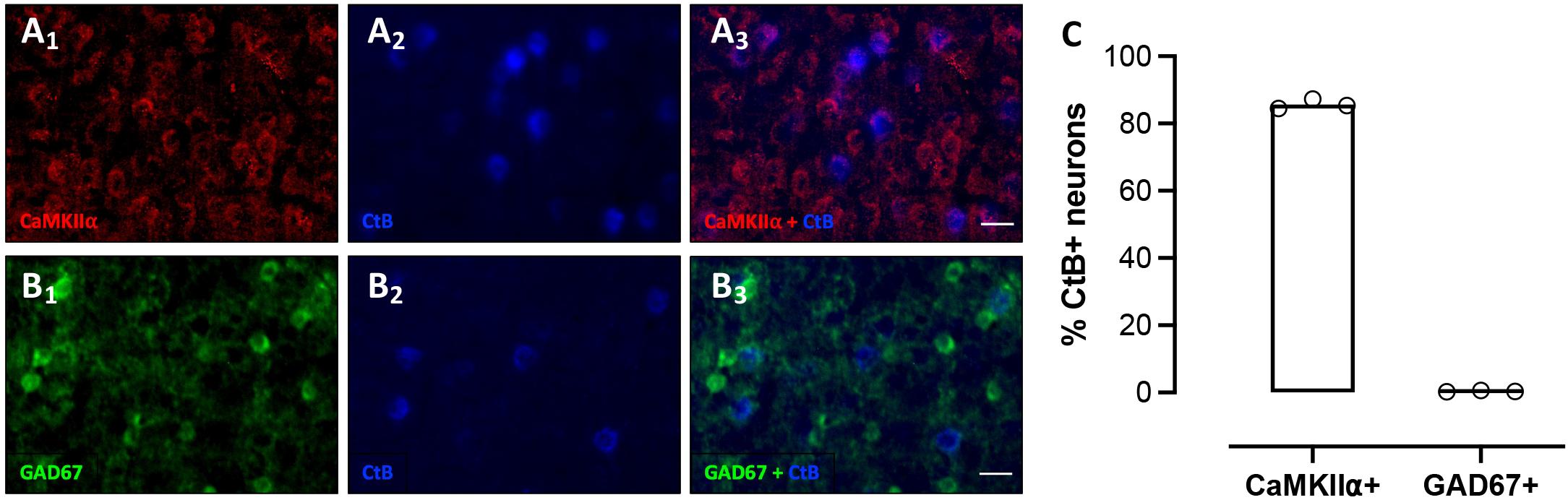
RMTg-projecting dmPFC neurons express CaMKIIα. Representative mPFC images co-labeled for **(A_1-3_)** the glutamatergic marker CaMKIIα (red) and CtB (blue) and the **(B_1-3_)** GABAergic marker GAD67 (green) and CtB (blue) from rat injected with CtB into the RMTg. **(C)** Quantification of co-labeling reveals that RMTg-projecting neurons are CaMKIIα^+^. Scale bar = 25 μm.

A number of cortico-subcortical projections purported to be involved in reward and aversion arise in layer V of the dmPFC. To examine whether RMTg-projecting dmPFC neurons may also overlap with populations of other subcortically projecting layer V neurons, neurons in this projection from four rats were selectively filled with green fluorescent protein using an intersectional, dual-virus approach (**Figure 3A**). Labeling was absent in one rat that was, therefore, excluded from analysis. In the remaining three rats, labeling was targeted to the dmPFC, was restricted to the injected hemisphere, and was not apparent in cell bodies outside of the dmPFC indicating successful isolation of the dmPFC-RMTg circuit (**Figure 3B**). Dense punctate labeling, indicative of synaptic terminals, was evident in a number of subcortical regions. Quantification of staining density relative to background (**Figure 3C-D**) revealed the greatest density of collaterals in the dorsomedial striatum (11.31 ± 3.11%), whereas the dorsolateral striatum was virtually devoid of labeling (0.07 ± 0.01%). Ventrally, RMTg-projecting dmPFC neurons collateralized to a moderate degree in the nucleus accumbens core (6.03 ± 1.31%) as well as the shell (0.34 ± 0.19%), albeit very weakly. Substantial collateralization was also observed in the ventral pallidum (11.05 ± 3.98%), hypothalamus (8.48 ± 0.98%), periaqueductal gray (6.54 ± 1.50%), and lateral preoptic nucleus (6.20 ± 0.74%). Terminal labeling was much less dense in the lateral habenula (5.15 ± 1.55%) and ventral tegmental area (2.58 ± 1.07%) – two regions heavily interconnected with both the dmPFC and the RMTg. By comparison, terminal labeling in the RMTg itself was 3.58 ± 0.17%. The amygdala also receives significant input from layer V dmPFC neurons and, like the RMTg, is well-known for its role in avoidance and aversive signaling. Despite this, only very weak terminal labeling was apparent in this region (0.94 ± 0.08%). The dorsal hippocampus (Hipp) and secondary somatosensory cortex (S2) were used as negative controls as it is well-known that these regions do not receive any input from the dmPFC (Hipp: 0.02 ± 0.01%; S2: 0.01 ± 0.00%).

**Figure 3.**
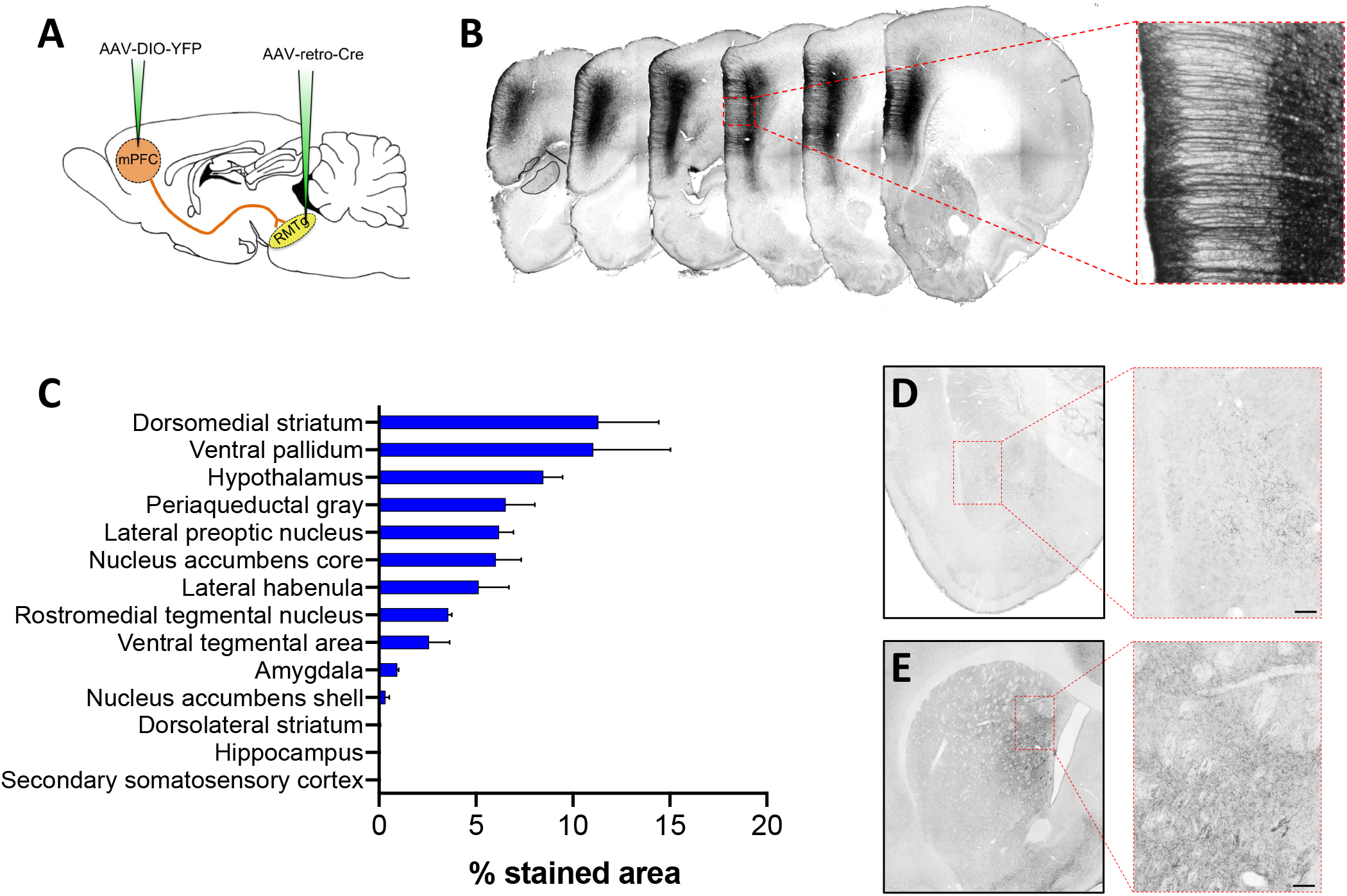
RMTg-projecting dmPFC neurons collateralize throughout the brain. **(A)** An intersectional dual-virus approach was used to fill RMTg-projecting dmPFC neurons with yellow fluorescent protein (YFP). **(B)** Representative images showing RMTg-projecting dmPFC neurons filled with YFP following amplification using standard immunohistochemistry. **(C)** Quantification of the average percent-stained area within ROIs placed within the respective brain regions. **(D)** Representative YFP staining in the amygdala shows relatively sparse collateralization of RMTg-projecting dmPFC neurons in the basolateral nucleus. **(E)** Representative YFP staining in the striatum shows dense collateralization in the dorsomedial but not dorsolateral striatum. Scale bar =100 μm.

### Selective stimulation of RMTg-projecting dmPFC neurons drives avoidance behavior

To begin to investigate whether dmPFC inputs to the RMTg play a significant role in aversive signaling, in-vivo optogenetics was used to measure real-time place preference in response to activation of this neural circuit. Testing on day 1 revealed that stimulation of dmPFC terminals in the RMTg resulted in significant avoidance of the light-paired compartment relative to chance (**Figure 4B**). This effect was replicated during testing on day 2 when the light-paired compartment was reversed [One-way ANOVA test 1 x test 2 x chance: F(1.95,7.75)=22.74; p=0.0006]. The magnitude of this avoidance was similar to that observed during stimulation of LHb terminals in the RMTg (**Figure 4C**), which was also significantly lower than chance [One-way ANOVA test 1 x test 2 x chance: F(1.59,7.95)=9.60; p=0.0095]. In contrast, stimulation of dmPFC terminals in the neighboring VTA resulted in neither preference nor avoidance of the light-paired compartment (**Figure 4D**) on either day 1 or day 2 [One-way ANOVA test 1 x test 2 x chance: F(1.04, 3.11)=0.095; p=0.7866]. Direct comparison of the effect of each circuit manipulation on real-time place preference revealed that stimulation of either dmPFC or LHb inputs to the RMTg drove avoidance behavior that was significantly different from stimulation of dmPFC inputs to the VTA (**Figure 4E**) [One-way ANOVA: F(2,12)=7.30, p=0.0084]. Altogether, these data indicate that, similar to the LHb-RMTg projection, activation of dmPFC inputs to the RMTg provides an aversive signal to promote avoidance behavior.

**Figure 4.**
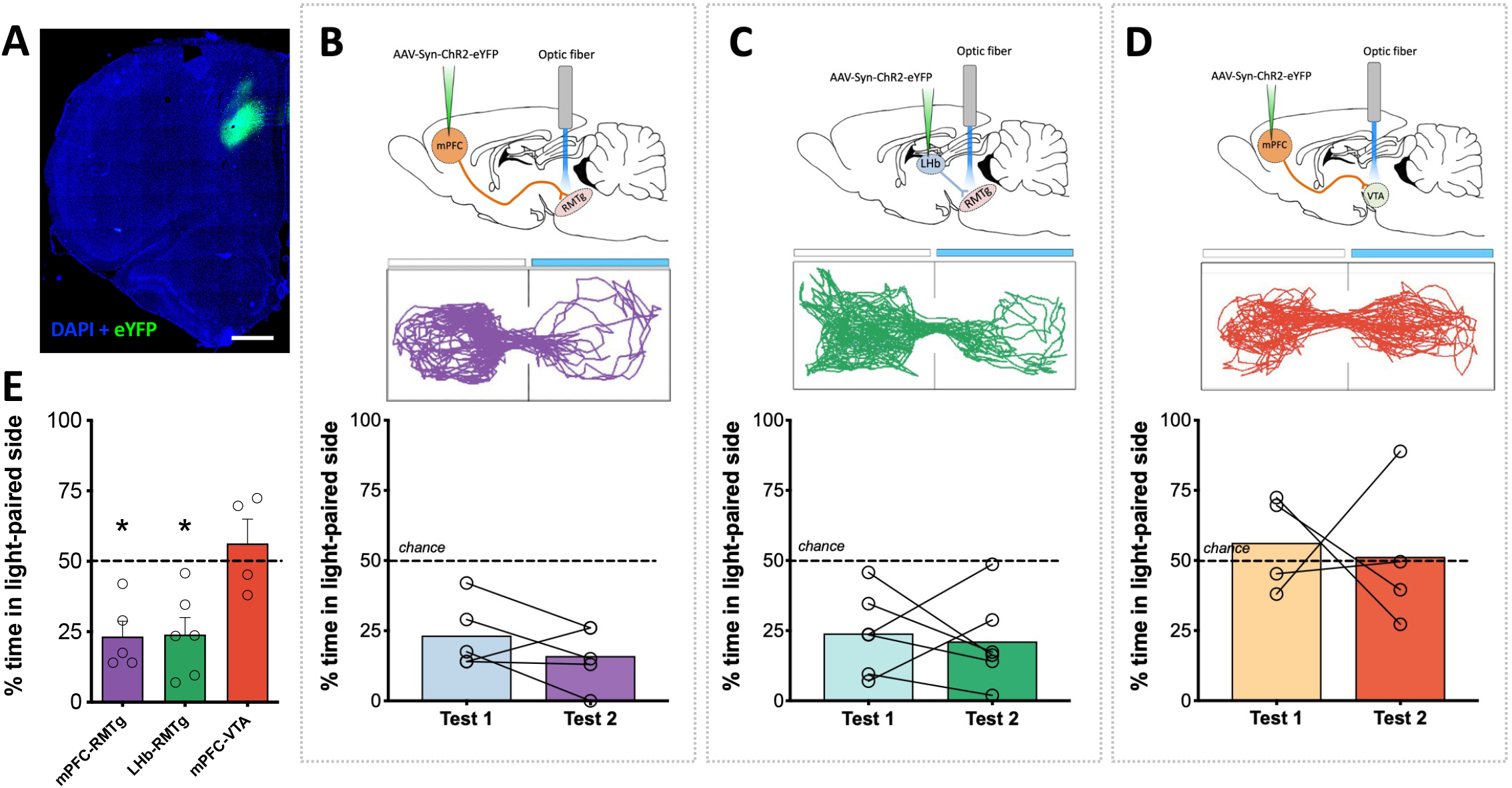
Optogenetic stimulation of RMTg-projecting dmPFC terminals drives avoidance. **(A)** Representative ChR2 expression in dmPFC. **(B)** Rats spend significantly less time relative to chance in the light-paired side of a two-compartment chamber during initial testing (test 1) and when the light-paired compartment is reversed (test 2) when light delivery results in stimulation of dmPFC terminals in the RMTg. **(C)** A similar degree of avoidance of the light-paired chamber is observed upon stimulation of lateral habenula inputs to the RMTg. **(D)** Unlike stimulation of dmPFC terminals in the RMTg, stimulation of dmPFC terminals in the VTA fails to produce either preference for or avoidance of the light-paired compartment. **(E)** Direct comparison of circuit manipulations reveals significant avoidance when stimulating inputs to the RMTg relative to the VTA. Light-paired side indicated by blue bar in representative maps above each dataset. *p≤0.01, scale bar = 1000 μm.

### RMTg-projecting mPFC neurons are activated following exposure to aversive stimuli

While optogenetic stimulation of dmPFC-RMTg neurons demonstrates that activation of this pathway can facilitate avoidance, we are unable to conclude from these data that neurons in this circuit are indeed active during avoidance and/or during similar behavioral responses to aversive stimuli. To explore this possibility, rats were euthanized 90 min following exposure to either neutral or aversive stimuli following injection of CtB into the RMTg (**Figure 5A**). Two groups of rats were exposed to a series of tones and foot shocks. In one group, tones were predictive of shock as in a standard fear conditioning paradigm. In contrast, in the other group, rats were exposed to the same number of tones and shocks but in an unpaired manner such that tones were not predictive of shock. As expected, rats in the Shock-paired tone group exhibited significantly greater freezing in response to tone presentation on test day than rats in the Shock-unpaired group (t-test; p=0.0005; **Figure 5B**). Behavioral data was not collected on the two remaining groups of rats exposed to either the neutral testing context or a series of foot shocks (without tone presentation). A one-way ANOVA comparing the magnitude of cFos expression in CtB^+^ neurons in the mPFC revealed a significant effect of treatment condition on cFos induction (**Figure 5C-D**) [F(3,32)=11.00, p<0.0001]. Post-hoc comparisons revealed that cFos expression was significantly greater in RMTg-projecting mPFC neurons of rats that were exposed to a series of either foot shocks or tones predictive of shocks relative to rats exposed to the neutral testing context (shock: p<0.0001; shock-paired tone: p=0.006). The magnitude of cFos expression in CtB^+^ mPFC neurons in shock-exposed rats was also significantly greater than was observed in rats exposed to tones that were explicitly unpaired with shocks (p=0.004). In combination with results from the real-time place preference testing, these data suggest that RMTg-projecting mPFC neurons are activated in response to learned and unlearned aversive stimuli and may play a role in regulating the behavioral response to such stimuli.

**Figure 5.**
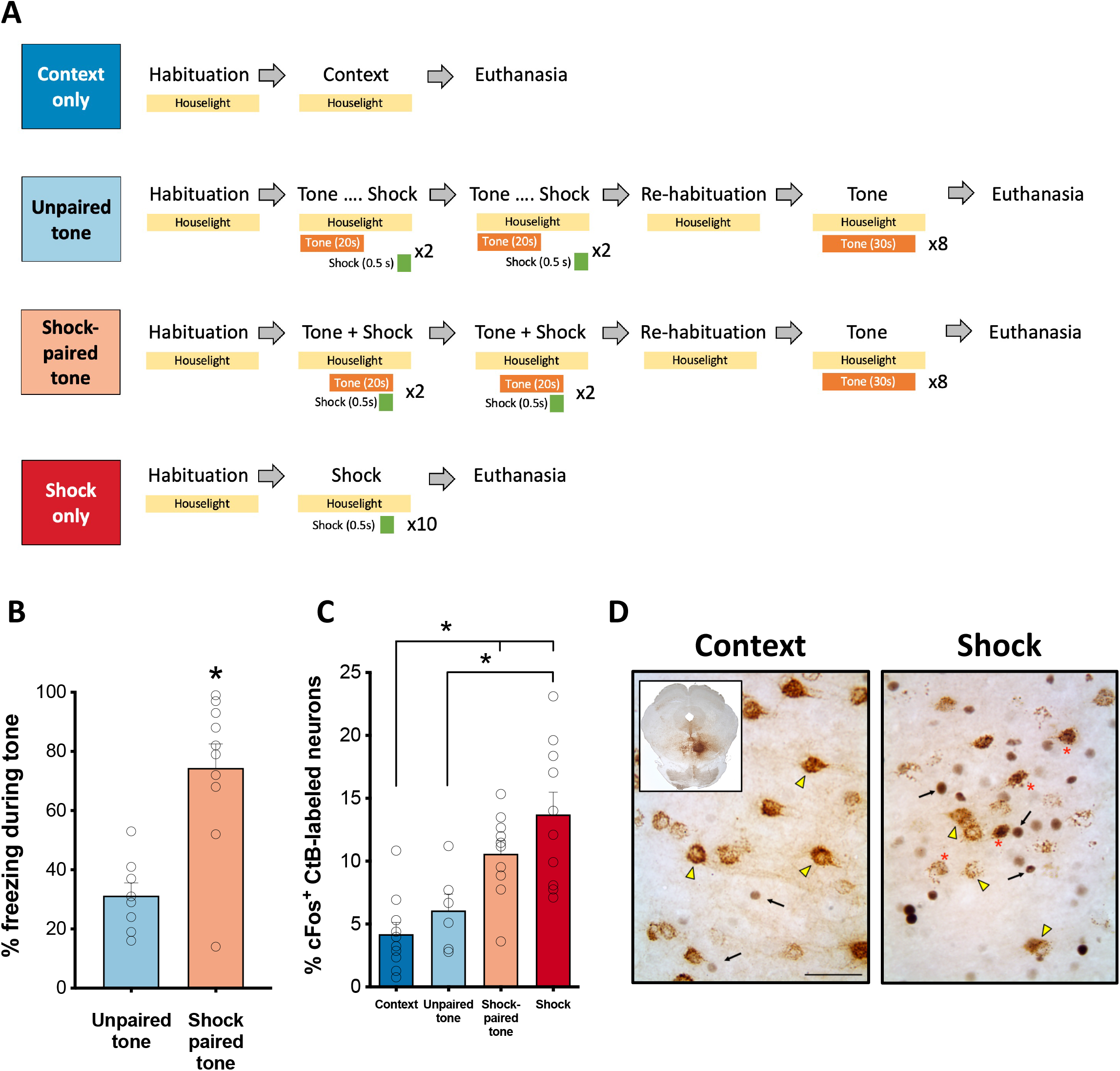
cFos induction in RMTg-projecting dmPFC neurons following exposure to aversive stimuli. **(A)** Experimental procedures. **(B)** Rats that had tone paired with foot shock delivery displayed significantly more freezing behavior in response to tone presentation than rats that were exposed to the same number of tone-shock presentations but in an unpaired manner. **(C)** Significantly greater cFos expression was observed in RMTg-projecting dmPFC neurons (CtB+) following exposure to either a series of foot shocks or a tone predictive of foot shock relative to a neutral tone or the testing context alone. **(D)** Representative images of CtB and cFos labeling in the dmPFC of a context-exposed rat and a rat exposed to foot shock. CtB^+^/cFos^-^ neurons are indicated with a yellow arrowhead; CtB^-^/cFos^+^ neurons are indicated with a black arrow; CtB^+^/cFos^+^ neurons are indicated by a red asterisk. Scale bar = 200 μm.

### Functional & structural changes in RMTg-projecting dmPFC neurons following exposure to aversive stimuli

Given the above evidence that dmPFC inputs to the RMTg are activated in response to aversive stimuli, we next investigated the potential impact that exposure to such stimuli has on plasticity in this neural circuit (**Figure 6A**). A two-way repeated measures ANOVA of spiking measured during whole-cell patch-clamp recordings from RMTg-projecting dmPFC neurons revealed a significant interaction between current step and stimulus exposure [F(15,315)=22.08, p<0.0001] such that spike frequency was significantly reduced as current injection increased beyond 200 pA in rats exposed to the same foot shock procedure that induced significant cFos expression relative to Context controls (Sidak correction; all p values ≤ 0.03; **Figures 6B-C**). This effect was accompanied by a significantly higher rheobase (t-test; p<0.0001), significantly lower membrane resistance (t-test; p<0.0001), and higher membrane capacitance (t-test; p<0.0434) in shock-exposed rats compared to context controls. No significant difference in action potential threshold was observed between groups (t-test; p=0.1555; **Figures 6D-G**). Action potential duration and amplitude were significantly different between groups with Shock-exposed rats exhibiting action potentials of greater amplitude (t-test; p=0.0232) and shorter duration (t-test; p=0.0002) than Context-exposed rats (**Figures 6H-I**). No significant difference in action potential afterhyperpolarization was observed between groups (t-test; p=0.2625; **Figure 6J**). Overall, these data are indicative of decreased intrinsic excitability in RMTg-projecting dmPFC neurons following exposure to an aversive stimulus.

**Figure 6.**
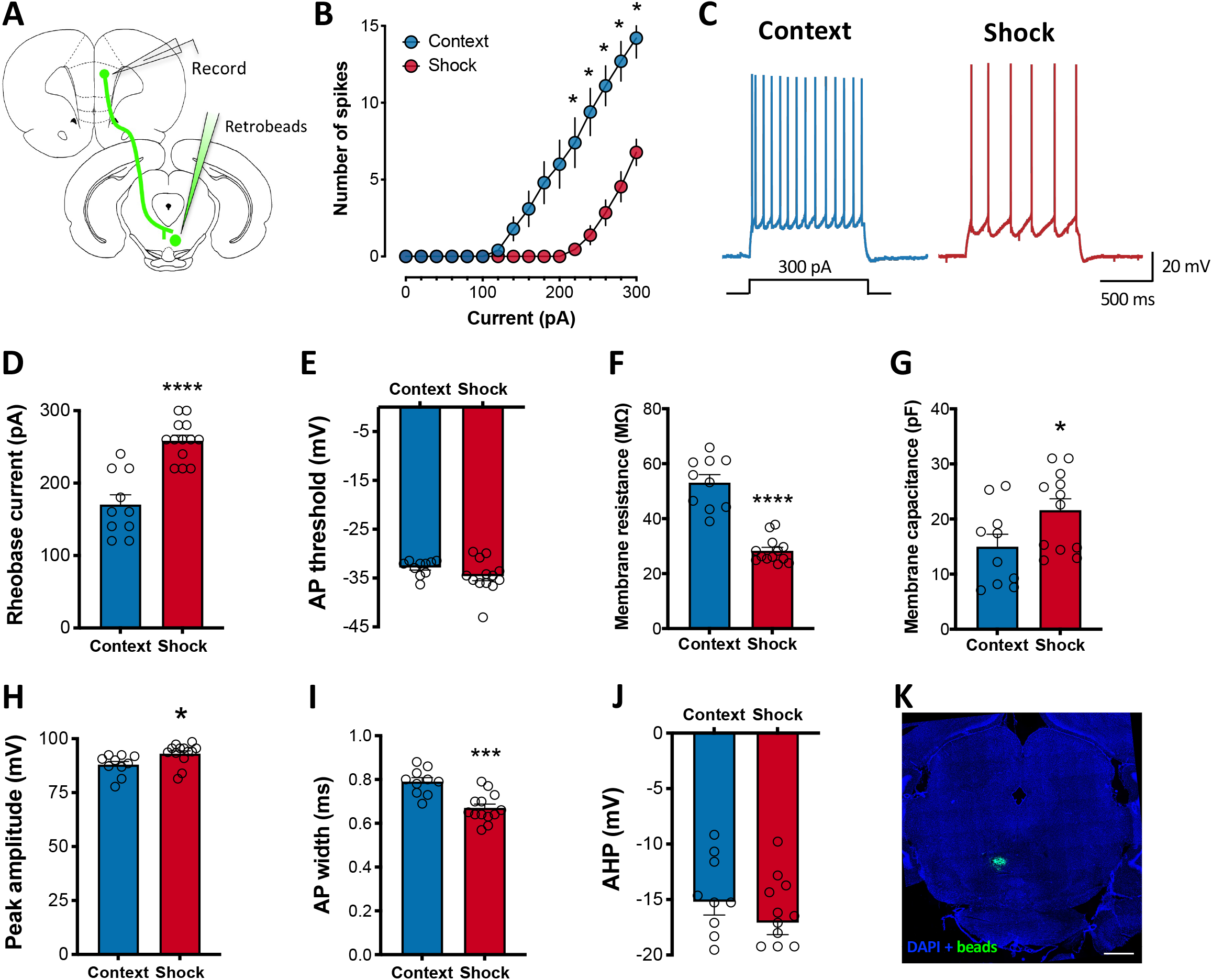
Decreased excitability in RMTg-projecting dmPFC neurons following exposure to aversive stimuli. **(A)** Experimental preparation. **(B)** Significantly fewer spikes were observed in shock-exposed rats relative to controls in current clamp recordings of retrobead-labeled dmPFC neurons. **(C)** Representative traces from a control and shock-exposed rat. Decreased spiking was associated with a significant increase in **(D)** rheobase, **(G)** membrane capacitance, and **(H)** peak action potential amplitude as well as a significant decrease in **(F)** membrane resistance and **(I)** action potential half-width. No significant difference was observed in **(E)** action potential threshold or **(J)** afterhyperpolarization. **(K)** Representative retrobead injection site in the RMTg. *p≤0.05, scale bar = 1000 μm.

To examine the impact of exposure to aversive stimuli on structural plasticity, we next quantified dendritic spine density and morphology in RMTg-projecting dmPFC neurons in rats exposed to either foot shock or a neutral context (**Figures 7A-B**). T-tests were used to analyze differences in dendrite diameter and volume as well as dendritic spine density collapsed across spine class. Two-way ANOVAs were used to analyze spine density and morphology by spine class between groups. No significant differences in dendrite diameter, volume or overall spine density were observed between Context- and Shock-exposed rats (*p* values > 0.50; **Table S1**). Analysis of spine density revealed a main effect of spine class [F(3,40)=53.37, p<0.0001] but no main effect of stimulus exposure [F(1,40)=0.075, p=0.7856] or interaction between the two factors [F(3,40)=1.34, p=0.2756]. Tukey corrected post-hoc comparisons of the main effect of spine class revealed that both Context- and Shock-exposed rats had a significantly greater density of mushroom-shaped spines relative to all other spine classes (all *p* values < 0.0001; **Figure 7C**). Of the eight other measures of spine morphology analyzed, no significant between-group differences were observed in dendritic spine length, diameter, or volume. Similarly, no significant between-group differences in spine terminal point diameter or volume were observed. Spine neck volume was not significantly different between groups either (**Table S1**). In contrast, a significant main effect of stimulus exposure [F(1,33)=9.85, p=0.0036] was observed for spine neck diameter in the absence of a significant interaction between stimulus exposure and spine class [F(3,33)=1.69, p=0.1884] indicative of greater spine neck diameter in Shock-exposed rats across all spine classes compared to Context-exposed rats (**Figure 7D**). A significant main effect of spine class was also uncovered with post-hoc analyses revealing that for both context- and shock-exposed rats, long, thin spines had significantly greater spine neck diameter than other spine classes (all p values ≤ 0.002). Differences in spine neck length were also observed between Context- and Shock-exposed rats (**Figure 7E**) with significant main effects of both spine class [F(3,29)=180.60, p<0.0001] and stimulus exposure [F(1,290)=4.19, p=0.0498]. While a significant interaction between the two factors was also uncovered [F(3,29)=3.18, p=0.0388], Sidak’s multiple comparisons failed to identify significant differences in spine neck length within any given spine class between Context- and Shock-exposed rats (all p values > 0.05).

**Figure 7.**
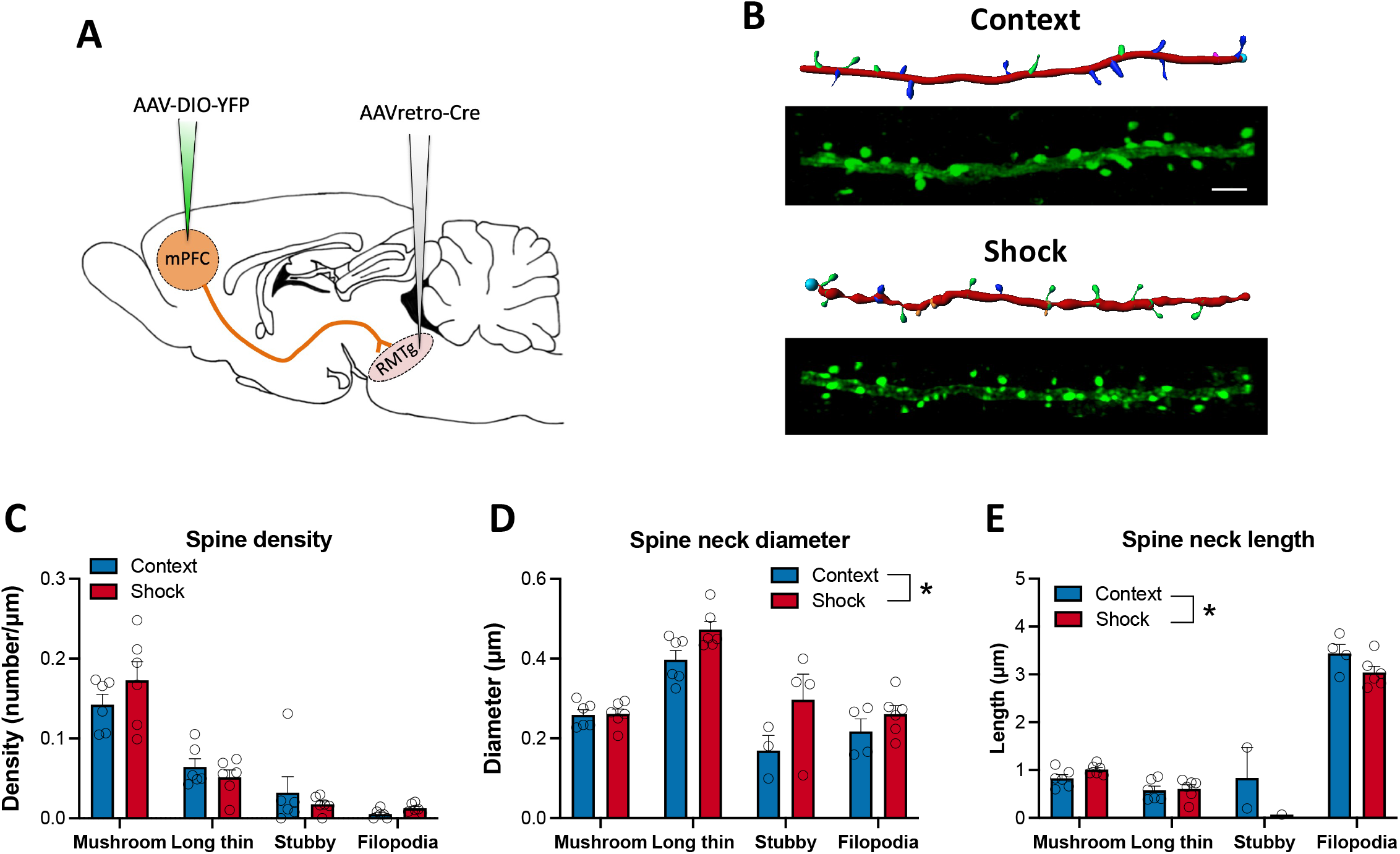
Exposure to aversive stimuli increases spine neck diameter in RMTg-projecting dmPFC neurons. **(A)** An intersectional dual-virus approach was used to fill RMTg-projecting dmPFC neurons with yellow fluorescent protein (YFP). **(B)** Representative YFP-filled primary apical dendrites in the dmPFC and accompanying IMARIS renderings for context- and shock-exposed rats. **(C)** Spine density did not differ between groups regardless of subclass. However, shock-exposed rats exhibited significantly greater spine neck diameter **(D)** and shorter spine length **(E)** across all subtypes (main effect of shock) relative to rats exposed to the neutral testing context. *p≤0.05; scale bar = 5 μm.

### RMTg-projecting dmPFC neurons express both D1 and D2 dopamine receptors

D1 and D2 dopamine receptors play important roles in prefrontal regulation of behavioral flexibility and decision-making (Floresco, 2013). A number of studies suggest that, similar to their distribution in the striatum, D1- and D2-expressing neurons in the mPFC are anatomically and functionally distinct cell populations (e.g., Gaspar et al., 1995). RNAScope for D1 and D2 receptor mRNA was used in combination with fluorescent retrograde tracing to investigate whether RMTg-projecting dmPFC neurons exhibit a distinct dopamine receptor expression profile and whether dopamine receptor gene expression was altered in these neurons following exposure to an aversive stimulus. As shown in **Figures 8A-F**, fluorescent retrobead-labeled cells (bead+) comprised approximately 62% of the total population of cells analyzed. A two-way ANOVA indicated that there was no significant difference in the number of either total or bead+ cells between control and shock-exposed rats [Main effect of treatment: F(1,12)=0.002, p=0.0966; treatment x cell-type: F(1,12)=0.10, p=0.759]. In addition, no significant between-group differences were observed using a two-way ANOVA to compare the percent of bead+ cells that were also positive for either D1 or D2 mRNA in control and shock-exposed rats [F(3,24)=0.28, p=0.8411] (**Figure 8G**). Collapsing across groups, classification of bead^+^ cells by dopamine receptor mRNA expression revealed that RMTg-projecting dmPFC neurons are predominantly D1 receptor-expressing (87%) with a substantial proportion also expressing D2 receptors (59%) (**Figure 8H**). Only a small proportion of bead+ neurons expressed D2 mRNA in the absence of D1 mRNA (5%) and approximately 8% lacked both mRNA for either receptor. To examine whether exposure to an aversive stimulus altered the magnitude of dopamine receptor gene expression in RMTg-projecting dmPFC neurons, two-way ANOVAs were used to compare the average number of RNAScope dots present per cell across dopamine receptor-expressing cell types. As shown in **Figures 8I-J**, foot shock exposure had no significant effect on either D1 (Treatment: F(1,18)=2.09, p=0.1651; treatment x cell-type: F(2,18)=0.21, p=0.8138) or D2 (Treatment: F(1,18)=1.38, p=0.2550; treatment x cell-type: F(2,18)=0.30, p=0.7445) mRNA expression.

**Figure 8.**
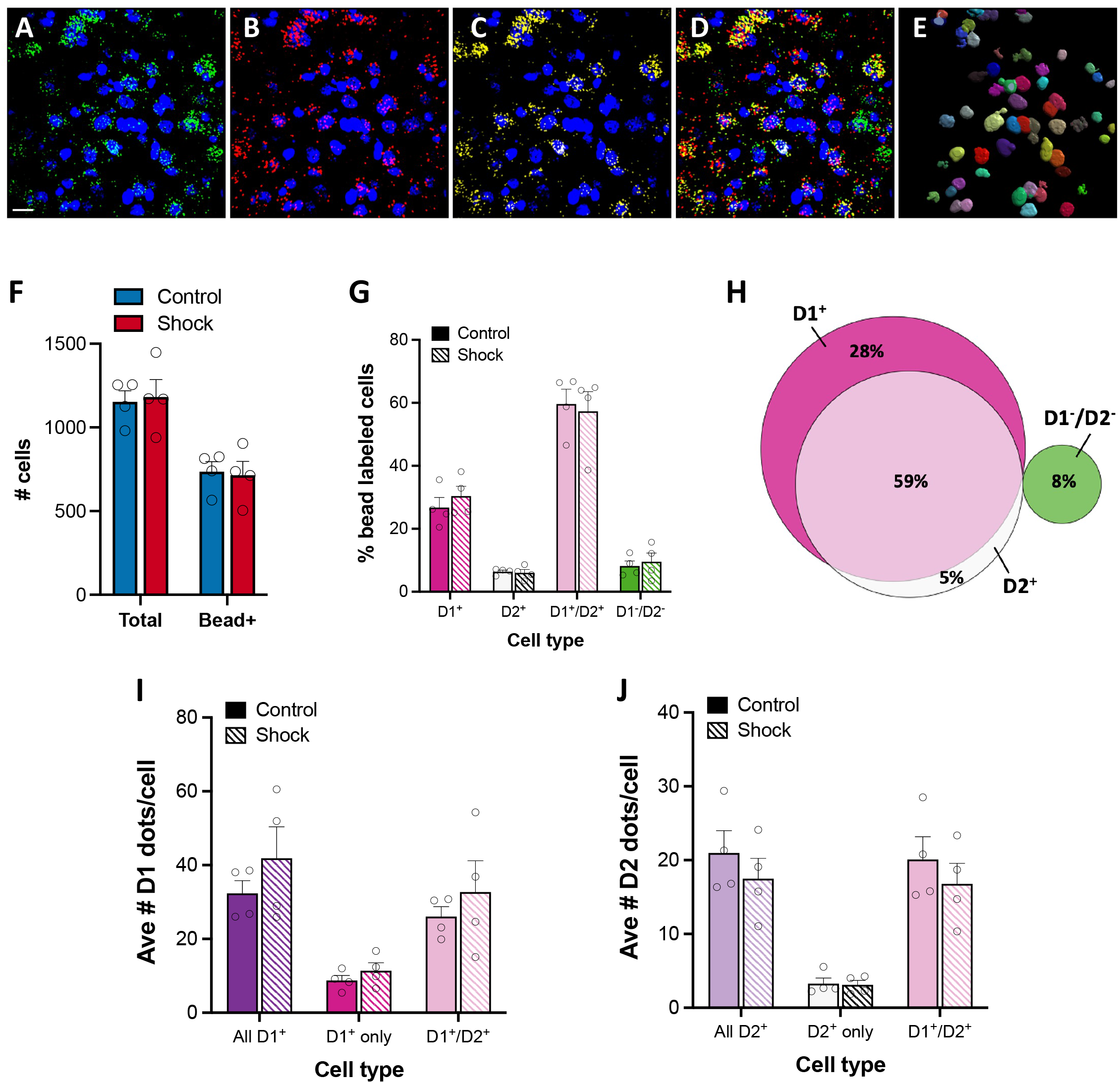
D1 & D2 receptor gene expression in RMTg-projecting dmPFC neurons. **(A-E)** Representative labeling in dmPFC neurons shows dense retrobead labeling in green **(A)** accompanied by D1 **(B)** and D2 **(C)** mRNA transcript dots indicated in red and yellow, respectively. The merged image **(D)** shows heterogeneous overlap of the retrograde beads and mRNA transcripts dots in cells identified using nuclear labeling with DAPI (blue). **(E)** IMARIS rendering of individual cells from A-D showing colocalization of the beads/dots in the 3D rendered soma. Foot shock had no effect on the total number of cells or number of bead positive cells analyzed **(F)**. The prevalence of D1 and D2 mRNA transcript containing cells was similar between the treatment groups **(G)**. When collapsed across groups **(H)**, it is apparent that the majority of RMTg-projecting dmPFC neurons are D1^+^ with a large proportion of D1^+^ neurons also expressing D2 receptor mRNA. Foot shock exposure had no significant effect on the magnitude of either D1 **(I)** or D2 **(J)** mRNA expression. Scale bar = 20 μm.

To further explore whether cFos induction observed following exposure to aversive stimuli is specific to a unique dopamine receptor-expressing population of dmPFC-RMTg neurons, we next measured induction of cfos mRNA colocalized with D1 receptor mRNA (the predominantly expressed dopamine receptor in these neurons). As expected, cfos expression was significantly enhanced in Shock-exposed rats relative to rats exposed to a neutral context (**Figure 9A**), and this was true regardless of retrobead labeling [two-way ANOVA main effect of treatment: F(1,12)=4.72, p=0.0505). When comparing cfos expression in D1^+^ and D1^-^ RMTg-projecting dmPFC neurons, a two-way ANOVA revealed a main effect of treatment with Shock-exposed rats exhibiting a greater proportion of cfos in both cell types relative to Context controls (**Figure 9B**). However, this effect only trended toward statistical significance for a main effect of shock exposure [F(1,12)=4.05, p=0.0673]. Similarly, the magnitude of cFos mRNA expression, as measured by average number of cFos dots per cell, was significantly greater in RMTg-projecting dmPFC neurons of Shock-exposed rats compared to controls regardless of D1 receptor expression profile (**Figure 9C**; main effect: F(1,12)=4.50, p=0.0555). Altogether, these data indicate that RMTg afferents arising in the dmPFC are highly enriched in D1 dopamine receptors (with D2 receptors colocalized to many of these neurons), and that the effects of exposure to aversive stimuli are similar across dmPFC-RMTg neurons with differing dopamine receptor expression profiles.

**Figure 9.**
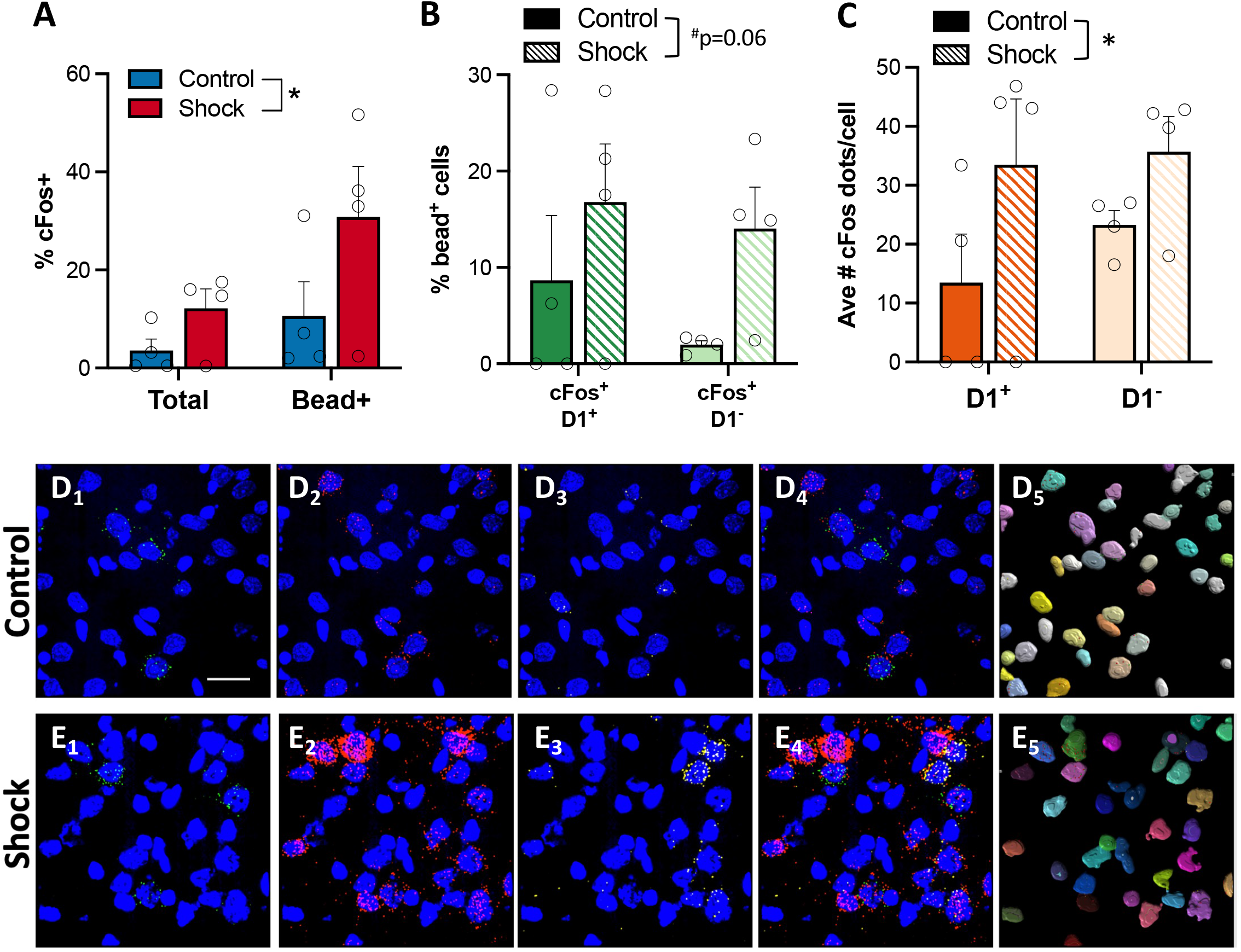
Shock-induced enhancement of cFos expression occurs in D1^+^ and D1^-^ RMTg-projecting dmPFC neurons. **(A)** Shock exposure significantly increased cFos mRNA expression in both bead^+^ and bead^-^ dmPFC neurons. cFos expression was similarly increased in both D1^+^ and D1^-^ neurons in shock-exposed rats relative to controls. **(C)** cFos mRNA expression was significantly greater in shock-exposed rats compared to controls, but but there was no significant difference between D1^+^ and D1 cell populations. Representative RNAScope labeling from a context-exposed control **(D)** and shock-exposed **(E)** rat. Retrobead labeling is depicted in green **(D-E_1_)**, cFos mRNA in red **(D-E_2_)**, D1 receptor mRNA in yellow **(D-E_3_)**, merged image **(D-E_4_)** and IMARIS rendering of colocalized beads and dots the cell soma **(D-E_5_)**. Nuclear labeling by DAPI depicted in blue. Scale bar = 20 μm; *p≤0.05.

## Discussion

Findings from the present study demonstrate the presence of dense cortical input to the RMTg spanning the entire rostrocaudal extent of the mPFC, OFC and AIC. The greatest density of afferents arises from the neurons in the dmPFC. The neurons are primarily D1^+^ with a large proportion also expressing D2 receptor mRNA. RMTg-projecting dmPFC neurons collateralize extensively in regions critically involved in regulating motivated behavior and flexible decision-making. Stimulation of dmPFC terminals in the RMTg drives avoidance and exposure to aversive stimuli induces cFos expression in this neural circuit. Finally, repeated exposure to an aversive stimulus results in significant alterations in RMTg-projecting dmPFC neurons in the form of increased excitability and changes in spine neck morphology. Together these data suggest that dmPFC neurons play an important role in governing the behavioral response to aversive stimuli.

If anatomical density is any indication of the influence a particular circuit may have over behavior, our data suggest that mPFC inputs to the RMTg are likely to play, at minimum, an equally important role in guiding adaptive responding to environmental stimuli as other heavily researched cortico-subcortical circuits. In a quantitative analysis of cortico-subcortical projection density in the mPFC, Gabbot et al. (2005) reported that ~8% of layer V PL and IL mPFC neurons project to the amygdala, whereas raphe- and PAG-projecting mPFC neurons each account for ~1-5% of neurons in layer V. Despite being sparse relative to PL and IL input to the ventral striatum (~18%), for example, subsequent work has implicated each of these discrete circuits in crucial aspects of motivated behavior (e.g., Rozeske et al., 2011; Warden et al., 2012; Bukalo et al., 2015). By comparison, the density of RMTg-projecting PL and IL neurons observed in the current study (~10%) is one of the denser subcortical projections arising from layer V of the mPFC. A large body of work has shown that the PL and IL mPFC exert opposing effects on many types of behavior. For example, PL mPFC neurons facilitate behavioral responding in Pavlovian and Instrumental assays associated with either appetitive or aversive outcomes, whereas IL mPFC activity has been shown to facilitate extinction of such behavioral responses (Peters et al., 2009; Gourley and Taylor, 2016). While the current study showed that stimulation of dmPFC terminals in the RMTg (which includes the PL subregion) facilitate avoidance, it remains unknown whether the IL-RMTg projection regulates RMTg signaling in an inverse manner that is similar to what is often observed when PL and IL mPFC are manipulated at a regional level. Unlike the PL and IL mPFC, the DP mPFC has been largely neglected with very few studies investigating this region at either anatomical or functional levels. In one of the few existing DP studies, Kataoka et al (2020) revealed a role for DP mPFC inputs to the dorsomedial hypothalamus in sympathetic stress response and stress-induced avoidance of social interactions. Combined with the current analysis, which reveals a remarkably dense projection from the DP mPFC to the RMTg that increases to an astonishing degree in the caudal mPFC, these data highlight the need for further investigation into the role of both the DP mPFC and its connections with the RMTg in aversion.

Consideration of collateral input is particularly important when investigating circuit function, as the possibility that manipulations of circuit activity affect signaling in sites that receive collaterals has the potential to influence interpretation of results. Using an intersectional, virally-mediated approach, our data reveal dense collateralization of RMTg-projecting dmPFC neurons to a number of regions critically involved in motivated behavior. While the extent of collateralization may be somewhat surprising, this frequently underappreciated aspect of neuronal structure is not uncommon. Indeed, recent methodological advancements have enabled researchers to map the extent of a single neuronal projection throughout the brain and demonstrate that collateralization is often widespread (Economo et al., 2016; Kebschull et al., 2016). The current findings indicate that RMTg-projecting dmPFC neurons collateralize most densely in the dorsal striatum. Interestingly, dmPFC afferents to the dorsal striatum are well-characterized for their role in guiding goal-directed behavior (Simmler and Ozawa, 2019) including avoidance (Loewke et al., 2021). Previous work found that, relative to a number of other cortico-subcortical projections, dmPFC input to the dorsal striatum was among the densest of projections, comprising ~19% of layer V neurons (Gabbott et al., 2005). Thus, it is not necessarily surprising that there is overlap in the population of dorsal striatum-projecting dmPFC neurons and those of other subcortical afferents. Unexpectedly, terminal density in the RMTg itself was relatively low by comparison to other brain regions. However, it should be noted that collaterals are often comprised of very thin branches (Rockland, 2013), and as a result the density measurement obtained in the present study (percent area stained) may not provide a full picture of the extent of collateralization of this projection. It is also unclear from the present data whether the observed collateralization was indicative of dense arborization of a select few dmPFC neurons, or of a high number of dmPFC cells each providing relatively weak collateral input to a given region. Recent work reporting very little overlap in cell body labeling between NAc- and RMTg-projecting mPFC neurons using dual retrograde tracer approach (Cruz et al., 2021) suggests that the former may be the more likely scenario. Additional experiments using multiple retrograde tracers to examine overlap in dmPFC cell body labeling will be essential to understand the potential functional implications of synergistic neurotransmission in regions receiving collateral input. Certainly, the current data present intriguing possibilities for coordinated signaling across brain regions involved in guiding behavioral responding to environmental stimuli.

The results of the present study also revealed that RMTg-projecting dmPFC neurons are glutamatergic and predominantly express D1 dopamine receptor mRNA. Of note, D2 receptor mRNA is also colocalized in the majority of these neurons. These findings agree in large part with existing data showing that D1 receptor expression is greater than that of D2 in the mPFC (Santana et al., 2009). While D1 and D2 receptor-expressing neurons are often thought of as discrete cell populations, a number of studies have observed colocalization of both receptors, particularly in layer V of the mPFC where D2 receptors are most abundant (Vincent et al., 1995; Gaspar et al., 1995; Santana et al., 2009). Recent work has highlighted the importance of dopaminergic regulation of cortical control in aversive signaling (Vander Weele et al., 2018; Huang et al., 2020). Of particular interest is data suggesting that dopamine signaling alters mPFC responses to aversive stimuli by altering the signal-to-noise ratio of incoming sensory inputs (Vander Weele et al., 2018). Whether this dopaminergic modulation is circuit- or cell-type specific is not well-understood. Nevertheless, a rich literature demonstrates that D1 and D2 receptors regulate behavioral flexibility in complex ways in the mPFC (Floresco and Magyar, 2006). The expression and function of both receptor subtypes is frequently mechanistically linked to neuropsychiatric illnesses characterized by deficits in decision-making including schizophrenia, addiction, and anxiety disorders (Volkow et al., 2004; Perez de la Mora et al., 2012; McCutcheon et al., 2019). While foot shock exposure did not significantly affect D1 or D2 mRNA expression in dmPFC-RMTg neurons in the current study, it is possible that repeated or prolonged insults uniquely affect dopamine receptor modulation of this neural circuit or that other measures of D1/D2 receptor function not explored in the current study are altered by exposure to aversive stimuli. Thus, the role that dopaminergic regulation of dmPFC-RMTg circuitry plays in aversive signaling and how it is altered in models of neuropsychiatric illness present intriguing areas for future investigation.

The present study focused on the potential role of dmPFC-RMTg circuit in aversion based on previous work characterizing the RMTg for its involvement in aversive signaling and the functional overlap exhibited by the dmPFC. However, some data suggests that PL-RMTg neurons respond to both aversive and rewarding stimuli (Li et al., 2019a). In addition, recent work suggests that activity in this neural circuit is particularly crucial when responding to conditioned rather than unconditioned stimuli (Li et al., 2019b; Cruz et al., 2020). It is therefore clear that additional work is needed to determine the potential diversity of signals encoded by these neurons and how this information differs across afferents arising from various mPFC subregions (i.e., PL vs IL).

Somewhat unexpectedly, exposure to repeated foot shock resulted in a significant decrease in excitability in RMTg-projecting dmPFC neurons. This contrasts with what has been observed in LHb neurons, the densest source of input to the RMTg (Jhou et al., 2009b), which exhibit a significant increase in excitability following a similar foot shock exposure paradigm (Lecca et al., 2016). Interestingly, the decrease in excitability observed in the present study was accompanied by significant changes in neck length and diameter of spines localized to the primary apical dendrites of RMTg-projecting dmPFC neurons. Dendritic spines are the primary recipients of incoming excitatory signals in pyramidal neurons. Spine neck morphology, in particular, plays a fundamental role in compartmentalizing electrical and biochemical signals in the head of the spine. In particular, spine neck diameter has been identified as the single greatest contributor to such compartmentalization (Tønnesen et al., 2014). In combination with data demonstrating an inverse relationship between spine neck diameter and excitatory potential (Araya et al., 2014), our data suggest a potential reduction in the synaptic strength of inputs to RMTg-projecting dmPFC neurons in shock-exposed rats relative to controls. Although speculative, given the role that layer V cortical apical dendrites are thought to play in modulating cortical oscillations (LaBerge and Kasevich, 2013), it is possible that the observed change in spine neck morphology alters oscillatory frequency in this neuronal population. Altogether, these data suggest a loss of top-down modulation of RMTg activity following exposure to aversive stimuli. Whether these physiological and structural adaptations serve to promote adaptive responding to future aversive stimuli is an important avenue for future exploration.

In conclusion, the current work presents a fresh perspective on the degree of subregion- and circuit-specific cortical regulation of RMTg-mediated aversive signaling. These data provide a strong foundation from which future studies can begin to dissect out the distinct (or complementary) roles of parallel cortico-subcortical circuits involved in motivated behavior. How these circuits are altered in models of neuropsychiatric illness will be crucial for understanding the neural mechanisms underlying disruptions in the balance of neural signals mediating reward and aversion that is altered in a number of disease states.

## Acknowledgements

The authors thank Joseph Pitock for technical support and Joroen Verharen for sharing the ImageJ cell density analysis protocol. We are also grateful to Sam Centanni for providing a detailed protocol for the analysis of the RNAScope data using Imaris Software.

---

**Table S1.** Complete analysis of dendritic spine density & morphology in RMTg-projecting dmPFC neurons following exposure to aversive stimuli. Bolded results indicate statistical significance of p≤0.05.

| Measure | Statistical test | Effect | Result |
| --- | --- | --- | --- |
| Dendrite diameter | Unpaired t-test |  | t(10)=0.6430, p=0.3960 |
| Dendrite volume | Unpaired t-test |  | t(10)=0.3731, p=0.1613 |
| Overall spine density | Unpaired t-test |  | t(10)=0.2997, p=0.4101 |
| Spine density by class | Two-way ANOVA | Class | **F(3,40)=53.37, p<0.0001** |
|  |  | Treatment | F(1,40)=0.0750, p=0.7856 |
|  |  | Class x Treatment | F(3,40)=1.338, p=0.2756 |
| Spine length by class | Two-way ANOVA | Class | **F(3,36)=224.7, p<0.0001** |
|  |  | Treatment | F(1,36)=1.427, p=0.2401 |
|  |  | Class x Treatment | F(3,36)=1.573, p=0.2129 |
| Spine diameter by class | Two-way ANOVA | Class | **F(3,36)=23.77, p<0.0001** |
|  |  | Treatment | F(1,36)=2.492, p=0.1232 |
|  |  | Class x Treatment | F(3,36)=0.8049, p=0.4994 |
| Terminal point diameter by class | Two-way ANOVA | Class | **F(3,36)=13.77, p<0.0001** |
|  |  | Treatment | F(1,36)=0.0579, p=0.8112 |
|  |  | Class x Treatment | F(3,36)=0.0967, p=0.9614 |
| Spine volume by class | Two-way ANOVA | Class | **F(3,36)=15.89, p<0.0001** |
|  |  | Treatment | F(1,36)=1.832, p=0.1843 |
|  |  | Class x Treatment | F(3,36)=0.7693, p=0.5188 |
| Spine neck volume by class | Two-way ANOVA | Class | **F(3,36)=18.86, p<0.0001** |
|  |  | Treatment | F(1,36)=2.404, p=0.1298 |
|  |  | Class x Treatment | F(3,36)=0.9704, p=0.4173 |
| Terminal point volume by class | Two-way ANOVA | Class | **F(3,36)=5.712, p=0.0026** |
|  |  | Treatment | F(1,36)=0.9216, p=0.3435 |
|  |  | Class x Treatment | F(3,36)=0.6379, p=0.5955 |
| Neck length by class | Two-way ANOVA | Class | **F(3,29)=180.6, p<0.0001** |
|  |  | Treatment | **F(1,29)=4.190, p=0.0498** |
|  |  | Class x Treatment | **F(3,29)=3.179, p=0.0388** |
| Neck diameter by class | Two-way ANOVA | Class | **F(3,33)=27.33, p<0.0001** |
|  |  | Treatment | **F(1,33)=9.847, p=0.0036** |
|  |  | Class x Treatment | F(3,33)=1.689, p=0.1884 |

